# Tau regulates epithelial morphogenesis through vesicle trafficking–dependent Notch activation

**DOI:** 10.64898/2026.03.23.713835

**Authors:** Neha Tiwari, Khushboo Sharma, Madhu G. Tapadia

**Author notes:** **Corresponding Author** – Madhu G. Tapadia; (Mob).

## Abstract

Tau is a conserved microtubule-associated protein best known for its roles in neuronal cytoskeletal stability and axonal transport. However, its functions in non-neuronal tissues remain poorly understood. Here we demonstrate that *Drosophila* Tau (dTau) regulates epithelial growth and tissue architecture in the *Drosophila* Malpighian tubules by controlling vesicular trafficking and Notch signaling. Loss of dTau results in pronounced epithelial hyperplasia, increased tubule diameter, and ectopic branching. Despite elevated Notch transcript levels, dTau-deficient tubules exhibit significantly reduced Notch intracellular domain (NICD), indicating impaired pathway activation. Proteomic and cellular analyses reveal widespread disruption of endocytic regulators and vesicle trafficking components, including reduced levels of the endocytic adaptor Liquid facets (Epsin) and altered distribution of Rab5, Rab7, and Rab11 endosomes. dTau loss also disrupts autophagic–lysosomal homeostasis and reduces endosome–lysosome fusion. These trafficking defects correlate with abnormal Delta localization and diminished Notch signaling. Together, our findings uncover a previously unrecognized role for dTau in maintaining epithelial signaling homeostasis by coordinating vesicular trafficking and receptor activation.

**Graphical Abstract:** 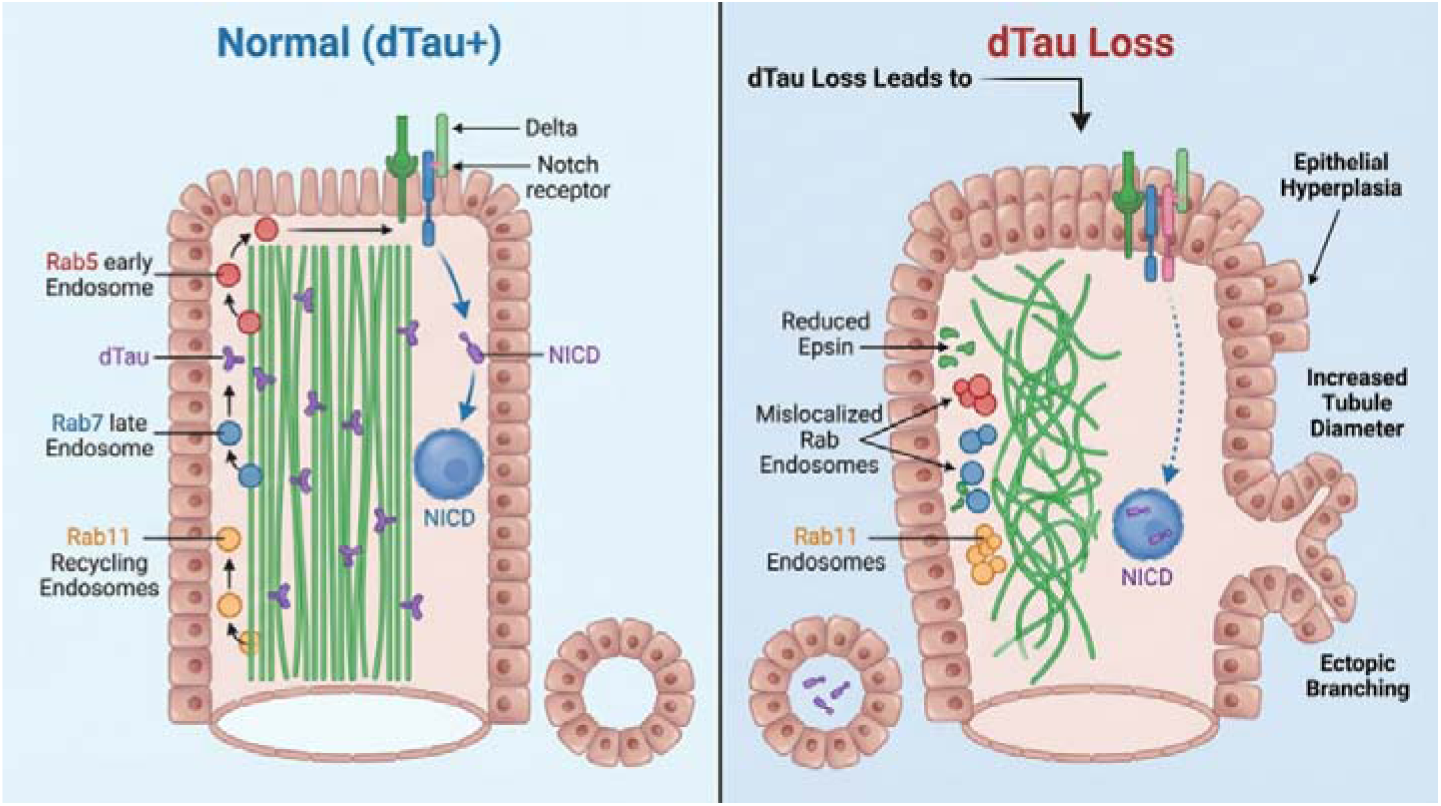

## INTRODUCTION

Tau is a conserved microtubule-associated protein best known for its neuronal roles in stabilizing microtubules in the neurons, regulating axonal transport, and maintaining cytoskeletal integrity (Frost et al., 2015, Goedert et al., 2019). Although Tau has been extensively studied in the context of neurodegeneration, emerging evidence indicates that Tau is also expressed in several non-neuronal tissues (Valles-Saiz et al., 2022; Lee et al., 2025; Tiwari and Tapadia, 2026), suggesting broader physiological functions that remain largely unexplored. Our recent work demonstrated that *Drosophila* Tau (dTau) plays a critical role in shaping the morphology of the *Drosophila* Malpighian tubules (MTs), a simple epithelial organ analogous to the vertebrate kidney. In the absence of dTau, MTs exhibit striking developmental defects, including cyst formation, increased tissue width, increased cell number, and early onset of morphological abnormalities beginning at embryonic stage 14. These defects are associated with altered cytoskeletal dynamics, increased Rho1 activity, destabilized microtubules, and disrupted epithelial convergence-extension movements. Despite such severe morphological abnormalities, dTau knockout (*tau ^KO^*) flies remain viable due to compensatory activity by the microtubule crosslinker Futsch, revealing a potential developmental buffering mechanism (Tiwari and Tapadia, 2026).

During these studies, we consistently observed excessive proliferation, ectopic branching, and tissue overgrowth in *tau ^KO^* MTs, hinting at a disruption in pathways governing epithelial growth control. The Notch signaling pathway is a highly conserved mechanism that regulates cell fate decisions across metazoans, with essential roles in *Drosophila* oogenesis, neurogenesis, myogenesis, and the development of the wing and eye (Fortini et al., 1993; Artavanis-Tsakonas et al., 1995; Egan et al., 1998; Kimble and Simpson, 1997). Canonical Notch activation begins when the Delta or Serrate ligand binds to the Notch receptor, triggering sequential proteolytic cleavages, including γ-secretase mediated release of the Notch intracellular domain (NICD), which then translocates to the nucleus to activate Notch-responsive genes involved in regulating proliferation, differentiation, and tissue morphogenesis (De Strooper et al., 1999; Mumm et al., 2000; Kopan and Ilagan, 2009). Its activation critically depends on tightly regulated intracellular trafficking of the receptor through endosomal compartments to facilitate efficient NICD production (Schnute et al, 2018; Zhou etal.2022; Rodriguez et al, 2023). In *Drosophila* MTs, Notch signaling contributes to early cell fate decisions and proliferation during tubule development, processes that ultimately influence the organization of principal and stellate cell populations (Duvall et al., 2022). These considerations prompted us to examine whether dTau contributes to MT growth and epithelial architecture by influencing Notch trafficking and pathway activation.

Here, we show that *tau ^KO^* tubules exhibit elevated cell proliferation, resulting in tissue overgrowth and branching phenotypesNotch levels are significantly reduced in *tau ^KO^* MTs. Proteomic analysis of *tau ^KO^* tubules identified differential expression of endocytic and vesicular trafficking pathways, including reduced levels of Liquid facets (Lqf/Epsin), a factor required for efficient Notch receptor activation and ligand internalization. Loss of dTau is accompanied by mislocalization of Rab5, Rab7, and Rab11 positive vesicles, accumulation of Ataxin2–positive RNA granules, and altered vesicle distribution, collectively consistent with impaired endosomal trafficking. In addition, markers of autophagic flux suggest perturbation of proteostatic pathways. These findings support a model in which dTau contributes to epithelial architecture and proliferation by maintaining vesicle trafficking integrity, preserving Notch signaling competence, and sustaining intracellular proteostatic balance.

## RESULTS

### Loss of dTau causes hyperplastic tubule growth and is associated with reduced Notch signalling

The *Drosophila* Malpighian tubules (MTs) form a simple epithelial excretory organ composed of paired anterior and posterior tubules extending from the gut (Fig. 1A-A’). In our previous study, we showed that loss of dTau causes pronounced morphological defects in MTs, including increased tubule diameter, ectopic branching, luminal cyst formation, and elevated numbers of principal cells (PCs) and stellate cells (SCs), indicative of epithelial hyperplasia. These abnormalities arise during late embryogenesis and persist throughout larval development, disrupting normal tubule architecture (Tiwari and Tapadia, 2026). Because epithelial growth and morphogenesis are often regulated by signaling pathways such as Notch (Wan et al., 2000; Nicolas et al., 2003; Bouras et al., 2008; Surendran et al., 2010), we next investigated whether the hyperplastic phenotype observed in *tau ^KO^*tubules is associated with altered Notch pathway activity

**Figure 1:**
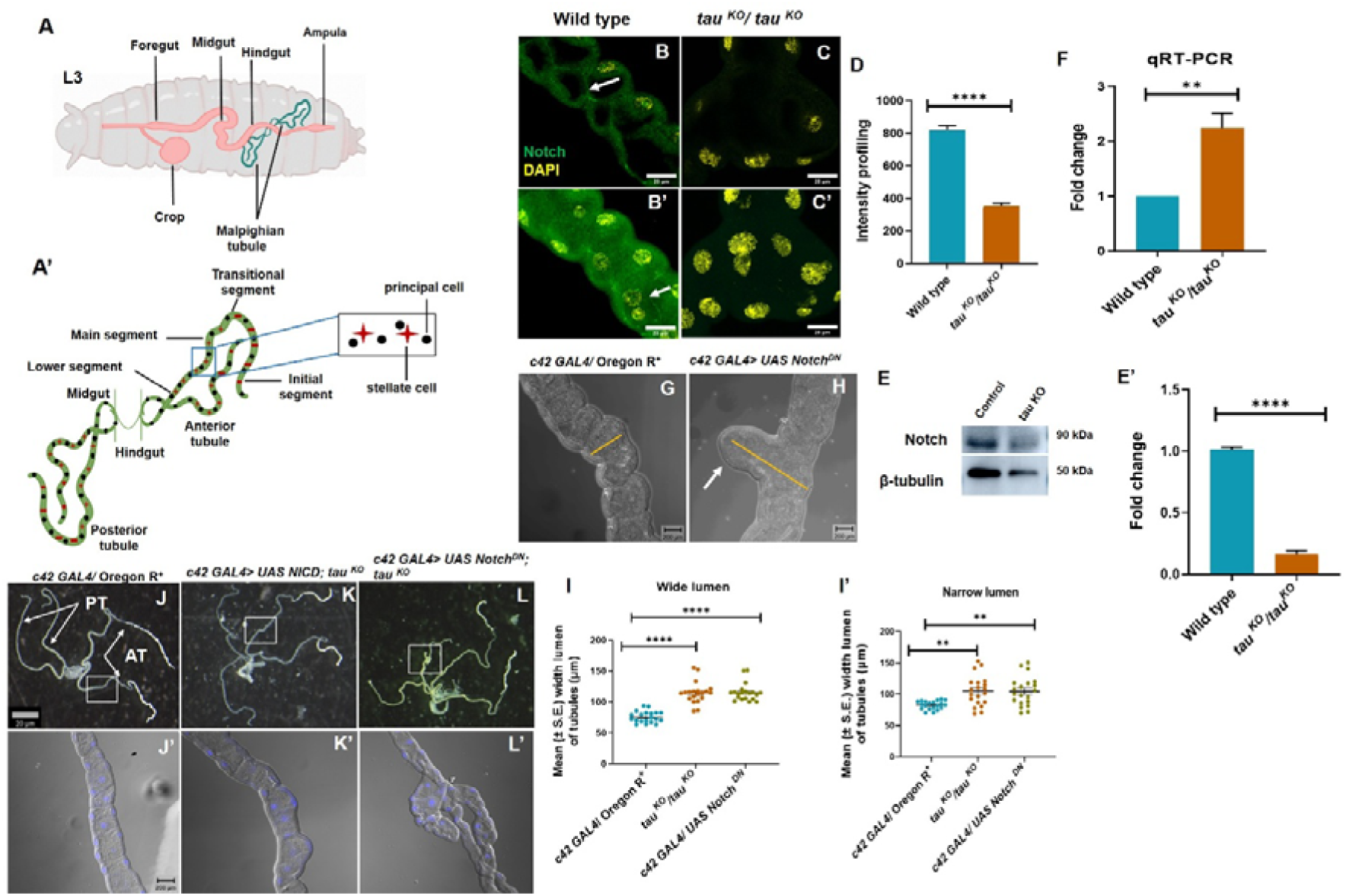
Loss of dTau reduces Notch signaling output and genetic modulation of Notch modifies *tau ^KO^*tubule phenotypes. **(A)** Diagrammatic representation of *Drosophila* Malpighian tubules (MTs). Larval gut (pink) with MTs shown in green. **(A’)** Classical morphology of larval MTs highlighting principal cells (PCs) and stellate cells (SCs). **(B, B**′) Wild-type tubules showing NICD (green) staining with DAPI (yellow). NICD signal is detectable in epithelial cells. **(C, C’)** *tau ^KO^* tubules showing NICD (green) staining. **(D)** Quantification of NICD fluorescence intensity in wild-type and *tau ^KO^* tubules (n=10). **(E-E’)** Western blot showing Notch protein levels. β-tubulin (M.W. 50 kDa) serves as a loading control (n=3). **(F)** RT–PCR analysis of Notch transcript levels in wild-type and *tau ^KO^* tubules (n=3). **(G-H)** DIC images showing Notch downregulation in a wild-type background (*c42-GAL4 > UAS-NotchDN*). **(I-I’)** Scatter dot plot showing the mean (±SE) lumen width, categorised as wide or narrow types (n = 20). **(J-L)** Bright-field images showing MTs from 3^rd^ instar larvae in different genetic background. **(J’-L’)** DIC images showing MTs from 3^rd^ instar larvae in different genetic background. All the Malpighian tubule images shown are from the wandering third-instar larvae. Scale bars represent 20 μm (B-C’) and 200 μm (F-G and I’-K’). All Images are maximum intensity projections of all the sections except (B and C) which are single section images. DAPI (yellow and blue)-stained nuclei. Statistical analysis was done using unpaired t-test and one-way ANOVA followed by Tukey’s multiple comparison test, **p < 0.01and ****p < 0.0001. Error bars, mean ± SE. All images represent 3 or more independent biological replicates.

To determine whether dTau influences Notch pathway activity in the MTs, we examined the localization and abundance of the Notch intracellular domain (NICD), the active form of the receptor. In wild-type third instar tubules, NICD was detected in epithelial cells, with prominent cytoplasmic signal (Fig. 1B, B’). In contrast, *tau ^KO^* tubules exhibited a marked reduction in overall NICD staining intensity (Fig. 1C, C’). Quantification of fluorescence intensity confirmed a significant decrease in NICD levels in *tau ^KO^* tubules compared to wild type (Fig. 1D). Consistent with the immunostaining results, western blot analysis of tubule lysates revealed a reduction in NICD protein levels in *tau ^KO^* samples relative to controls (Fig. 1E), and densitometric quantification confirmed a significant decrease after normalization to loading control (Fig. 1E’). To assess whether this reduction is a consequence of altered transcription, we measured Notch mRNA levels. RT–PCR analysis revealed increased Notch transcript levels in *tau ^KO^* tubules compared to control (Fig. 1F). These findings suggest that in the absence of dTau there is enhanced expression of Notch mRNA levels and, in spite of enhanced Notch transcription, reduced NICD levels were observed. These findings support the hypothesis that impaired Notch signaling could be contributing to the hyperplastic and morphological defects associated with Tau loss.

To directly validate this hypothesis, we genetically manipulated Notch signaling in the MTs following the system of Brand and Perrimon,1993. Using MTs specific *c42-GAL4* driver (Rosay et al., 1997), we downregulated Notch activity by a dominant-negative *UAS-Notch-DN* construct. It was observed that *c42-GAL4 > UAS-Notch-DN* in an otherwise wild-type background resulted in an increased tubule diameter, epithelial disorganization, and cyst-like enlargements (Fig. 1G–I’). These abnormalities recapitulated several key morphological features observed in *tau ^KO^*tubules, including increased PCs and SCs cell numbers in condition (Fig. S1D-E). Decreased NICD staining and reduced Notch transcript levels confirmed the efficacy of *c42 GAL4> UAS UAS-Notch-DN* (Fig. S1A–C′). We next asked whether restoring Notch activity could modify the *tau ^KO^* phenotype. Expression of constitutively active NICD in the *tau ^KO^* background (*c42-GAL4 > UAS-NICD; tau ^KO^*) partially rescued tubule morphology. Reduction in excessive branching and epithelial disorganization (Fig. 2K, K’) was observed, along with a decrease in the total cell number relative to *tau ^KO^* tubules (Fig. S1F, G). Conversely, further reduction of Notch signaling in the *tau ^KO^* background (*c42-GAL4 > UAS-Notch-DN; tau ^KO^*) markedly exacerbated the phenotype, resulting in severely thickened and highly branched tubules with pronounced epithelial irregularities (Fig. 1L, L’).

**Figure 2:**
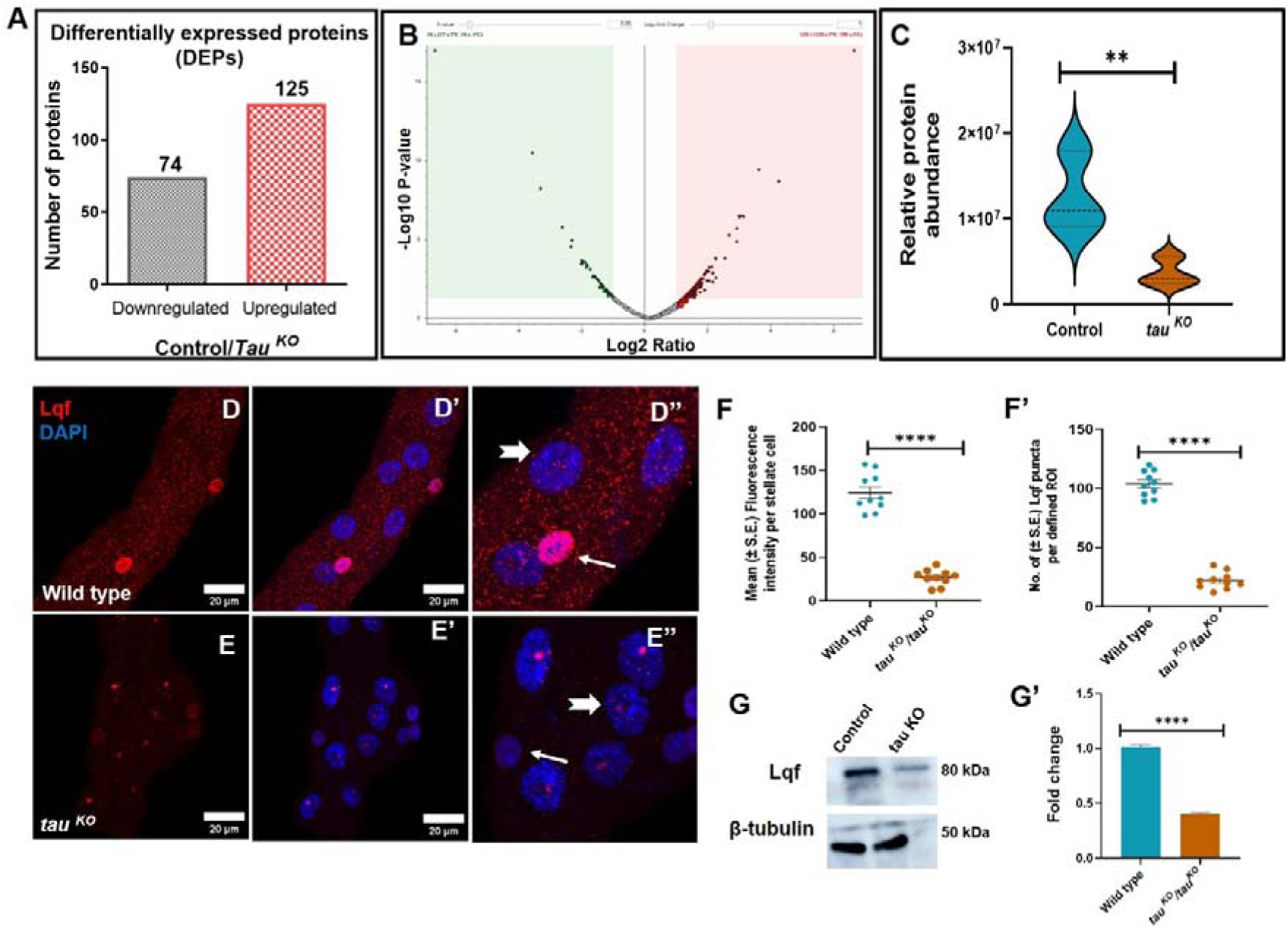
Loss of Tau disrupts endocytic regulators required for Notch activation in MTs. **(A)** Bar graph showing the number of significantly upregulated and downregulated proteins in *tau ^KO^* samples compared to control. **(B)** Volcano plot representation of the proteomic dataset showing statistical significance versus fold change of proteins detected in *tau ^KO^* tubules relative to control. **(C)** Violin plot showing normalized relative protein abundance of Liquid facets (Lqf) in control and *tau ^KO^* MTs based on proteomic analysis across biological replicates (n=3). **(D–D**″) Immunostaining of Lqf (red) in wild-type MTs. **(E–E**″) Immunostaining of Lqf (red) in *tau ^KO^* MTs. **(F)** Scatter plot showing quantification of Lqf fluorescence intensity per stellate cell in wild-type and *tau ^KO^* tubules (n=10). **(F**′) Scatter dot plot showing quantification of Lqf-positive puncta within a defined region of interest (ROI) in wild-type and *tau ^KO^* tubules (n=10). **(G)** Western blot analysis of Lqf protein levels in control and *tau ^KO^* MTs. β-tubulin (M.W. 50 kDa) serves as a loading control. **(G**′) Densitometric quantification of western blot bands showing relative Lqf protein levels normalized to loading control(n=3). All the Malpighian tubule images shown are from the wandering third-instar larvae. All images are maximum intensity projections of all the sections. DAPI (blue)-stained nuclei. Scale bar represents 20 μm. Statistical analysis was done using unpaired t-test, ****p < 0.0001. Error bars, mean ± SE. All images represent 3 or more independent biological replicates.

Together, these gain- and loss-of-function analyses indicate that dTau is required to regulate appropriate Notch signaling output in the MTs. Reduced Notch activity contributes to the growth and patterning defects observed in *tau ^KO^* tubules, and partial restoration of Notch signaling alleviates key aspects of the Tau-dependent phenotype.

### Proteomic analysis reveals disruption of endocytic regulators required for Notch activation in Tau-deficient tubules

Given that Notch transcript levels were elevated while NICD protein levels were reduced in *tau ^KO^* tubules, we hypothesized that dTau regulates Notch signaling post-transcriptionally. To identify molecular pathways affected by dTau depletion, we performed comparative proteomic analysis of wild-type and *tau ^KO^* Malpighian tubules. Global proteomic profiling revealed widespread changes in protein abundance between the two conditions. In total, 74 proteins were significantly downregulated and 125 proteins were upregulated in *tau ^KO^* tubules relative to controls (Fig. 2A). Volcano plot analysis highlighted a subset of proteins displaying significant fold change and statistical significance (Fig. 2B). Functional categorization of differentially expressed proteins indicated that many of the downregulated proteins were associated with vesicle trafficking, endocytosis, and cytoskeletal organization, whereas upregulated proteins were enriched for metabolic and stress-associated functions. The proportional distribution of functional categories among upregulated and downregulated proteins is summarized in supplementary pie-chart analyses (Fig. S2 A, B). Thus, it is possible that defects in intracellular trafficking is necessary for receptor-ligand interaction.

Among the downregulated candidates, the endocytic adaptor Liquid facets (Lqf/Epsin) was of particular interest because of its well-established role in Delta ligand internalization and productive activation of the Notch signaling pathway (Overstreet et al., 2003; Wang and Struhl, 2004). Quantitative proteomic measurements revealed a significant reduction in Lqf levels in *tau ^KO^* tubules compared with wild type, as illustrated by violin plot representation of normalized protein abundance values across biological replicates (Fig. 2C). To validate this proteomic observation, we examined Lqf localization by immunostaining. In wild-type Malpighian tubules, Lqf displayed a punctate cytoplasmic distribution (Fig. 2D–D’’). Higher-resolution analysis revealed cell-type–specific differences in Lqf localization: stellate cells showed prominent nuclear enrichment of Lqf (arrow), whereas principal cells primarily exhibited cytoplasmic punctate staining (arrowhead). In contrast, *tau ^KO^* tubules exhibited a significant reduction in overall Lqf signal intensity together with a decrease in cytoplasmic puncta and reduced stellate-cell nuclear enrichment (Fig. 2E–E’’). Measurement of fluorescence intensity per stellate cell demonstrated a significant reduction in Lqf signal in *tau ^KO^* tubules relative to controls (Fig. 2F), and analysis of puncta density within defined regions of interest revealed a corresponding decrease in Lqf-positive vesicular structures (Fig. 2F’). Consistent with the immunostaining results, western blot analysis of tubule lysates showed reduced Lqf protein levels in *tau ^KO^* samples, with densitometric quantification confirming a significant decrease compared with wild type after normalization to loading controls (Fig. 2G, G’).

To examine whether Lqf levels influence Notch pathway activity in MTs, we manipulated Lqf expression using the *c42-GAL4* driver. RNAi-mediated knockdown of Lqf resulted in a clear reduction of NICD staining, indicating diminished Notch signaling (Fig. S2D–D″). Conversely, overexpression of Lqf restored NICD signal comparable to control tubules (*c42 GAL4*/ Oregon R^+^) (Fig. S2E–E″), and intensity profiling showed comparable NICD fluorescence in control and *c42 GAL4> UAS Lqf* condition relative to the knockdown condition (Fig. S2F). To verify the efficiency of these genetic manipulations, RT–PCR analysis was performed, which confirmed reduced Lqf transcript levels in the RNAi line and elevated Lqf transcript levels in the overexpression condition relative to controls (Fig. S2G). These results indicate that Lqf dosage influences Notch pathway activation in the MTs epithelium.

Because efficient Notch signaling critically depends on Delta internalization and its progression through the endocytic pathway, the reduction of Lqf in *tau ^KO^* tubules suggested that Delta localization and trafficking might be disrupted in the absence of dTau.

### Tau is required for proper Delta trafficking in Malpighian tubule cells

Because Lqf plays a central role in Delta internalization during Notch activation, we next examined whether loss of dTau affects the localization of Delta, the Notch ligand, in Malpighian tubules. In wild-type tubules, Delta was detected both at the plasma membrane and within discrete cytoplasmic puncta, consistent with its dynamic trafficking through the endocytic pathway (Fig. 3A, A’). Notably, Delta signal appeared relatively well-organized around SCs nuclei.

**Figure 3:**
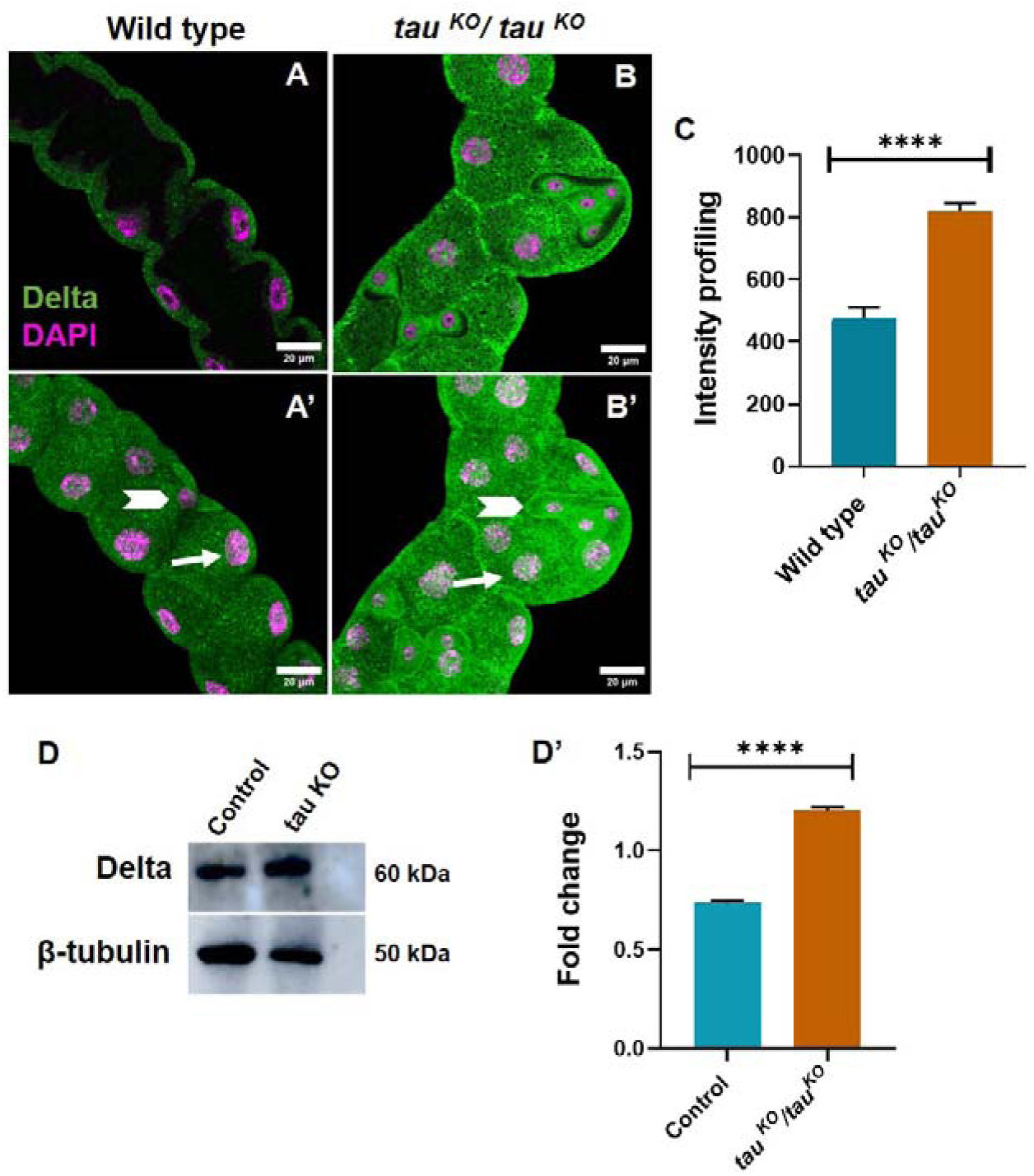
Tau loss disrupts intracellular distribution of the Notch ligand Delta in MTs. (A, **A**′) Immunostaining of Delta (green) in wild-type third instar MTs. Delta (green) is detected at the plasma membrane and in discrete cytoplasmic puncta. **(B, B**′) Immunostaining of Delta in *tau ^KO^* Malpighian tubules. **(C)** Quantification of cytoplasmic Delta fluorescence intensity in wild-type and *tau ^KO^* MTs showing a significant increase in cytoplasmic Delta signal in *tau ^KO^* tubules (n=10). **(D)** Western blot analysis of Delta protein levels in control and *tau ^KO^* MTs. β-tubulin (M.W. 50 kDa) serves as a loading control. **(D**′) Densitometric quantification of Delta protein levels normalized to loading control, showing increased Delta accumulation in *tau ^KO^* tubules (n=3). All the Malpighian tubule images shown are from the wandering third-instar larvae. All images are maximum intensity projections of all the sections except (A and B) which are single section images. DAPI (magenta) stained nuclei. Scale bar represents 20 μm. Statistical analysis was done using unpaired t-test, ****p < 0.0001. Error bars, mean ± SE. All images represent 3 or more independent biological replicates.

In contrast, *tau ^KO^* tubules displayed a pronounced alteration in Delta distribution. Instead of the punctate intracellular pattern observed in controls, Delta exhibited strong membrane enrichment together with extensive cytoplasmic accumulation (Fig. 3B, B’), suggesting impaired trafficking or altered intracellular handling of the ligand. This altered distribution was also evident around SCs nuclei, where Delta signal appeared more and less organized compared to wild type. Quantitative analysis confirmed a significant increase in cytoplasmic Delta signal in *tau ^KO^* tubules compared with wild type (Fig. 3C).

To further examine whether dTau loss affects overall Delta protein levels, we performed western blot analysis of MTs lysates, which revealed elevated Delta protein levels in *tau ^KO^* tubules, consistent with the increased cytoplasmic accumulation observed by immunostaining. (Fig. 3D, D’).

Together, these findings indicate that dTau contributes to the proper intracellular distribution of Delta in MTs cells. Given the known requirement of Delta endocytosis for productive Notch activation, disruption of Delta trafficking in *tau ^KO^* tubules may contribute to the reduced Notch signaling observed in the absence of dTau.

### Loss of Tau disrupts endosomal organization and vesicle trafficking pathways

To assess whether dTau loss broadly affects vesicular transport, we examined the distribution of Rab GTPases that define distinct endosomal compartments. In wild-type MTs, Rab5, a marker of early endosomes, localized to small and discrete cytoplasmic puncta characteristic of early endosomal structures (Fig. 4A–A’’). In contrast, *tau ^KO^* tubules exhibited enlarged and clustered Rab5-positive structures (Fig. 4B–B’’), indicating altered early endosome organization. Quantitative analysis revealed a significant increase in the average size of Rab5-positive puncta in *tau ^KO^* tubules compared to wild type (Fig. 4C). In addition, puncta number was quantified by counting Rab5-positive vesicles within a defined region of interest (ROI), which showed a significant increase in puncta density in *tau ^KO^* tubules relative to wild type (Fig. 4C’). These results suggest accumulation and enlargement of early endosomal compartments upon dTau loss.

**Figure 4.**
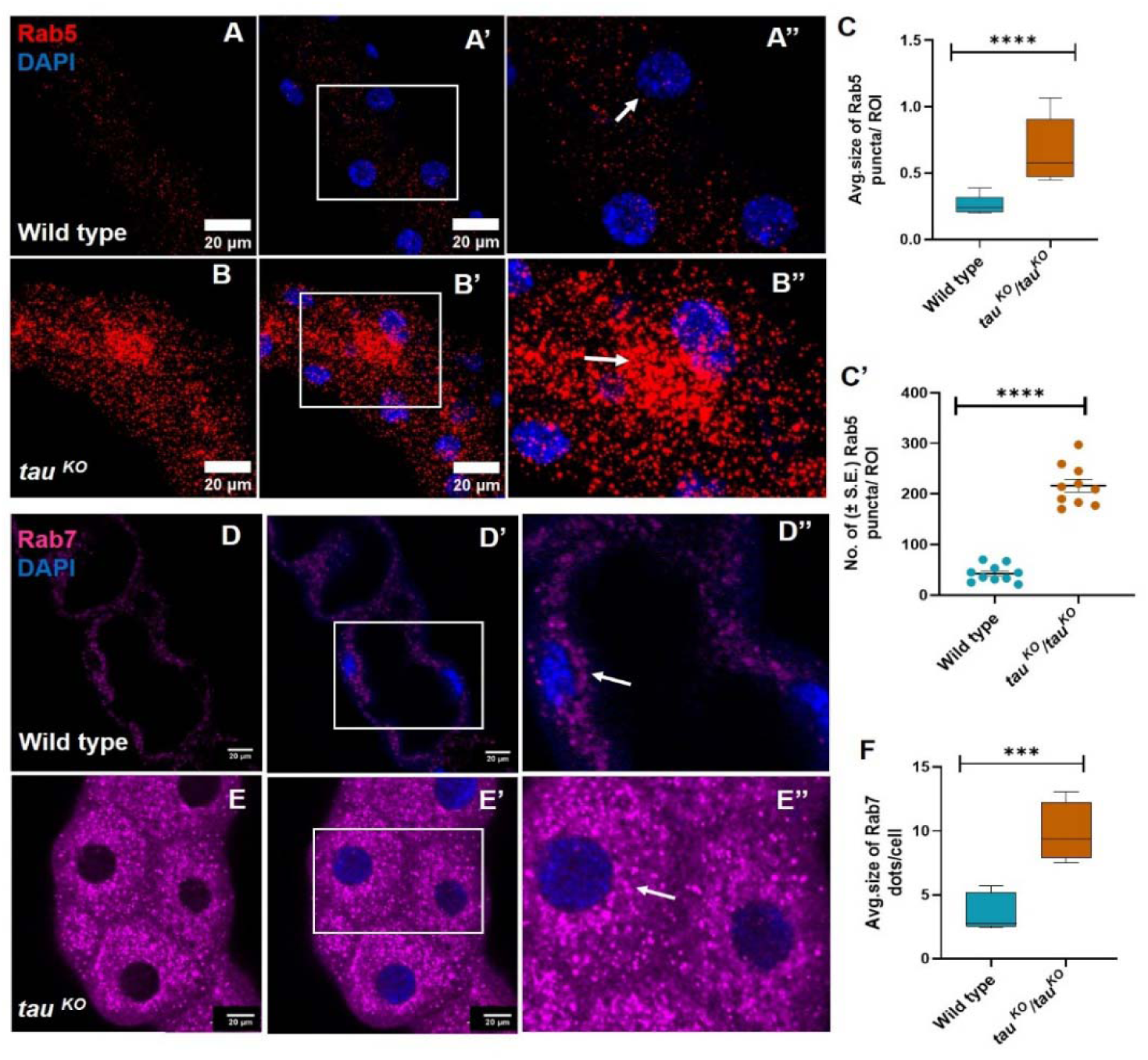
Tau loss disrupts endosomal organization in MTs cells. **(A–A**″) Immunostaining of Rab5 (red) in wild-type MTs showing small, discrete cytoplasmic puncta. The boxed region in A’ is shown at higher magnification in A″. Arrow indicates representative Rab5-positive vesicles. **(B–B**″) Immunostaining of Rab5 (red) in *tau ^KO^* MTs. Boxed regions are shown at higher magnification in B″. Arrow indicates representative enlarged Rab5-positive vesicles. **(C)** Box and whisker plot showing quantification of the average size of Rab5-positive puncta in wild-type and tau *^KO^* tubules (n=10). **(C**′) Scatter dot plot showing quantification of Rab5 puncta number per defined region of interest (ROI) showing increased vesicle density in tau *^KO^* tubules compared with wild type (n=10). **(D–D**″) Immunostaining of Rab7 (magenta) in wild-type MTs showing punctate vesicles. Boxed regions are shown at higher magnification in D″. Arrow indicates representative Rab7-positive vesicles. **(E–E**″) Immunostaining of Rab7 in *tau ^KO^* MTs. Boxed regions are shown at higher magnification in E″. Arrow indicates representative Rab7-positive vesicles. **(F)** Box and whisker plot showing quantification of average Rab7 puncta size in wild-type and *tau ^KO^* tubules (n=10). All the Malpighian tubule images shown are from the wandering third-instar larvae. All images are maximum intensity projections of all the sections except (D-E’) which are single section images. DAPI (blue) stained nuclei. Scale bar represents 20 μm. Statistical analysis was done using unpaired t-test, ***p<0.001 and ****p < 0.0001. Error bars, mean ± SE. All images represent 3 or more independent biological replicates.

We next examined Rab7, which labels late endosomes. In wild-type tubules, Rab7 displayed a punctate cytoplasmic distribution corresponding to late endosomal vesicles (Fig. 4D–D’’). In *tau ^KO^* tubules, however, Rab7-positive structures appeared enlarged and densely clustered (Fig. 4E–E’’). This altered Rab7 distribution suggests disruption of late endosomal organization and impaired progression through the endocytic pathway. Furthermore, quantitative analysis showed increase in the average size of Rab7 dots per cell (Fig. 4F)

We next examined Rab11, a marker of recycling endosomes. In wild-type tubules, Rab11 localized to well-defined punctate structures within the cytoplasm, consistent with active recycling compartments (Fig. S3A-A’). In contrast, Rab11 staining in *tau ^KO^* tubules was markedly reduced and poorly organized, suggesting defective recycling endosome function and impaired receptor recycling (Fig. S3B-B”). Quantification of Rab11 fluorescence intensity revealed a significant reduction in *tau ^KO^*tubules compared with wild type (Fig. S3C).

In addition to alterations in Rab-defined endosomal compartments, we observed changes in the localization of Ataxin-2, a component of ribonucleoprotein granules that has been linked to vesicle trafficking and cellular stress responses (Satterfield and Pallanck, 2006; Nonhoff et al., 2007). In wild-type tubules, Ataxin-2 displayed a punctate cytoplasmic distribution and was largely excluded from the tubule lumen (Fig. S3D–D′). In contrast, *tau ^KO^*tubules exhibited prominent Ataxin-2–positive aggregates, with dense material accumulating within the tubule lumen (Fig. S3E–E′). Quantitative analysis revealed a significant increase in the average size of Ataxin-2–positive puncta per cell in *tau ^KO^* tubules compared with wild-type controls (Fig. S3F). These structures may reflect altered ribonucleoprotein organization and are consistent with broader disruptions in intracellular trafficking and cellular homeostasis upon dTau loss.

Together, these findings indicate that dTau contributes to the proper organization of early, late, and recycling endosomal compartments in MTs cells. Loss of dTau leads to accumulation of Rab5- and Rab7-positive structures and disruption of Rab11-defined recycling endosomes, suggesting impaired endosomal trafficking that may contribute to the defects in Delta trafficking and Notch signaling observed in *tau ^KO^* tubules.

### Tau deficiency impairs autophagic and lysosomal function in Malpighian tubules

Given the accumulation of Rab5- and Rab7-positive endosomal vesicles in *tau ^KO^* MTs, we next examined whether loss of dTau affects autophagic–lysosomal degradation pathways, which are essential for maintaining cellular homeostasis and organelle turnover. To assess autophagic activity, we first examined the localization of Atg8, a core component of autophagosome membranes. In wild-type tubules, Atg8 displayed a sparse punctate distribution within the cytoplasm, consistent with basal autophagic activity (Fig. 5A–A’). In contrast, *tau ^KO^* tubules showed a marked reduction in Atg8-positive puncta (Fig. 5B–B’). Quantification of Atg8 puncta revealed a significant decrease in autophagosomal structures in *tau ^KO^* tubules compared with controls (Fig. S4A), indicating reduced autophagic activity. To further monitor autophagosome formation, we analyzed a *3XAtg8-mCherry* reporter, which labels autophagic vesicles in vivo. Consistent with the Atg8 immunostaining results, control tubules displayed numerous punctate autophagic structures (Fig. 5C-C’), whereas *tau ^KO^* tubules exhibited fewer and less prominent Atg8-mCherry–positive vesicles (Fig. 5D-D’), supporting the conclusion that loss of dTau compromises autophagosome formation and autophagic activity in Malpighian tubules.

**Figure 5.**
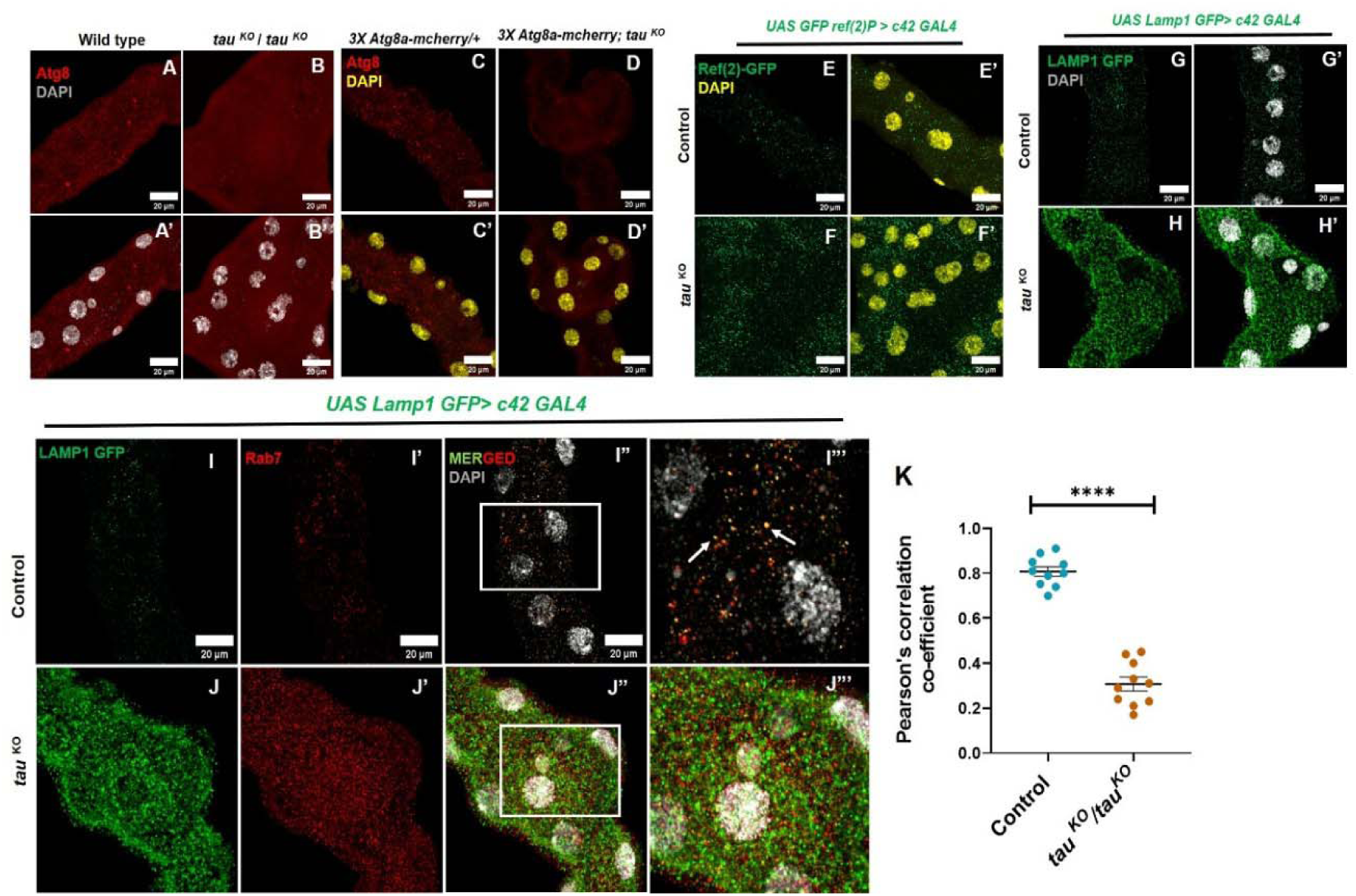
Loss of Tau disrupts autophagy and lysosomal organization in *Drosophila* MTs. **(A–A′)** Immunostaining for Atg8 (red), a marker of autophagosomes, in wild type MTs **(B-B’)** Immunostaining for Atg8 (red) in *tau ^KO^* tubules, exhibit a marked reduction in Atg8-positive puncta. **(C–C**′) Visualization of autophagosomes using the *3×Atg8-mCherry* reporter. Control tubules show abundant Atg8-positive autophagic structures. **(D-D’)** Visualization of autophagosomes using the *3×Atg8-mCherry* reporter in *tau ^KO^*tubules, displayed fewer and fewer autophagic vesicles. **(E–E**′) Analysis of autophagic cargo using *Ref(2)P-GFP*. In control tubules Ref(2)P signal is low and diffusely distributed. **(F-F’)** Analysis of autophagic cargo using *Ref(2)P-GFP*. *tau ^KO^* tubules exhibited prominent accumulation of Ref(2)P-positive aggregates. **(G–G**′) Lysosomal organization visualized using *Lamp1-GFP*. Control tubules display small punctate lysosomes distributed throughout the cytoplasm, **(H-H’)** Lysosomal organization visualized using *Lamp1-GFP*. In *tau ^KO^* tubules shows enlarged and clustered Lamp1-positive structures. **(I–I**D) Colocalization analysis of Lamp1-GFP (green) and Rab7 (red) in control tubules reveals frequent overlap between late endosomes and lysosomes. Insets show magnified regions highlighting Rab7–Lamp1 puncta (arrows). **(J–J**D) Colocalization analysis in *tau ^KO^* tubules. Rab7-positive vesicles accumulate and show reduced colocalization with Lamp1-positive lysosomes. Insets show magnified regions highlighting Rab7–Lamp1 puncta. **(K)** Colocalization between Rab7 and Lamp1 was quantified using Pearson’s correlation coefficient using FIJI JACoP plugin. All the Malpighian tubule images shown are from the wandering third-instar larvae. All images are maximum intensity projections of all the sections. DAPI (grey and yellow) stained nuclei. Scale bar represents 20 μm. Statistical analysis was done using unpaired t-test, ****p < 0.0001. Error bars, mean ± SE. All images represent 3 or more independent biological replicates.

**Figure 6:**
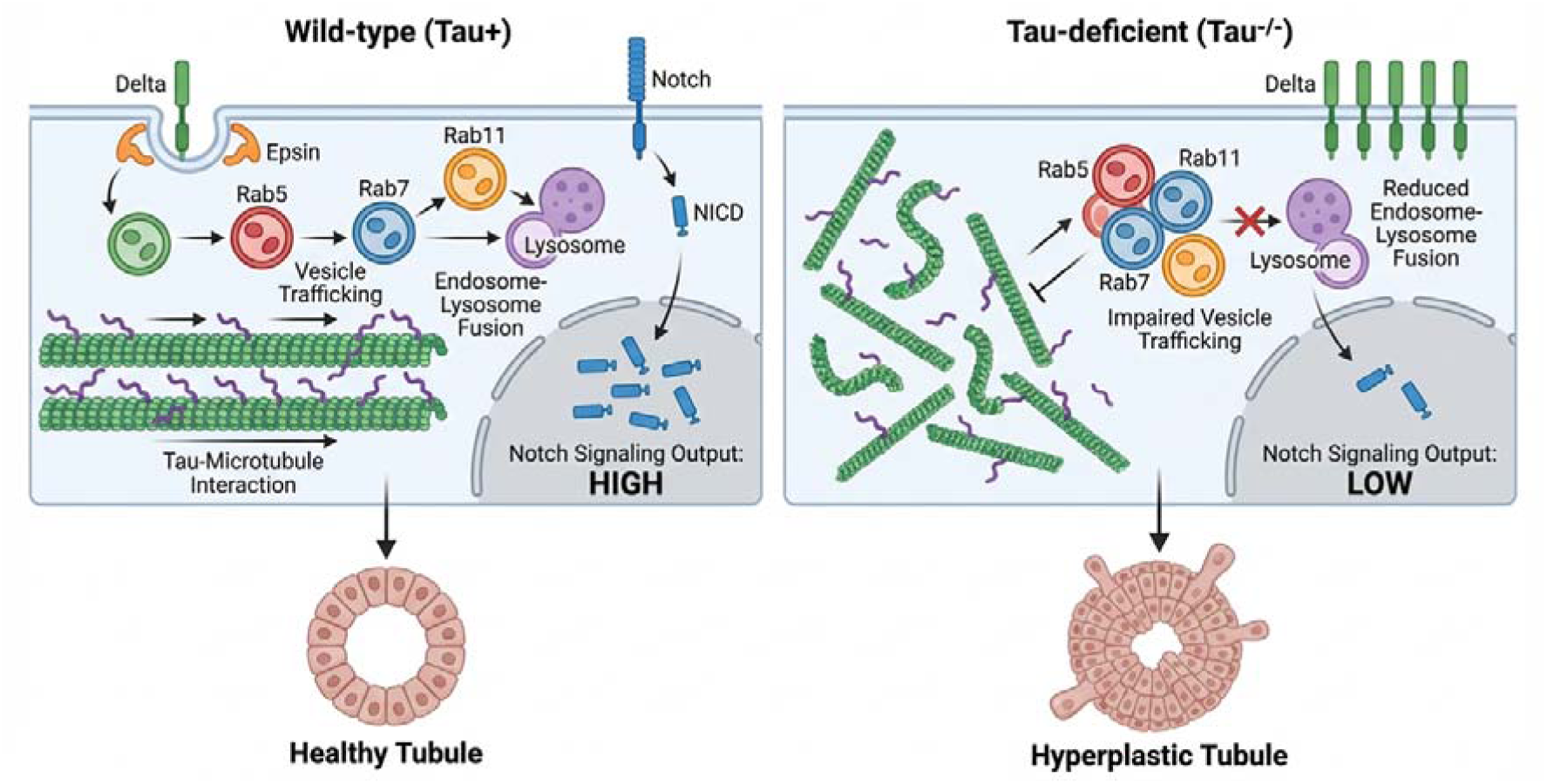
Proposed model illustrating the role of Tau in regulating vesicle trafficking and Notch signaling during Malpighian tubule morphogenesis. In wild-type MTs, Tau stabilizes microtubules and supports efficient vesicle trafficking through the endosomal pathway. Proper organization of Rab5-, Rab7-, and Rab11-positive compartments facilitates Delta trafficking and Notch receptor activation, leading to generation of the Notch intracellular domain (NICD) and balanced epithelial proliferation. Autophagic–lysosomal homeostasis is maintained, as indicated by normal Atg8 puncta, low Ref(2)P accumulation, and efficient Rab7–Lamp1 fusion. In *tau ^KO^* tubules, loss of Tau disrupts microtubule-dependent vesicle transport, leading to altered endosomal organization, defective Delta trafficking, and reduced Notch signaling. These defects are accompanied by impaired autophagic activity, accumulation of Ref(2)P aggregates, and altered lysosomal organization. Together, these alterations result in epithelial hyperplasia, branching defects, and abnormal tubule morphology.

We next investigated whether autophagic cargo degradation was affected using *Ref(2)P-GFP*, the *Drosophila* homolog of the autophagy adaptor p62, which accumulates when autophagy is impaired. In control, *Ref(2)P-GFP* signal was low and diffusely distributed, consistent with efficient cargo degradation (Fig. 5E–E’). In contrast, *tau ^KO^* tubules exhibited prominent accumulation of Ref(2)P-positive aggregates (Fig. 5F–F’). Quantification confirmed a significant increase in Ref(2)P puncta in *tau ^KO^* tubules (Fig. S4B), indicating impaired autophagic cargo clearance. To determine whether lysosomal organization was also affected, we examined the distribution of Lamp1-GFP, a marker for lysosomal compartments. In control tubules, *Lamp1-GFP* labeled small punctate lysosomes evenly distributed throughout the cytoplasm (Fig. 5G-G’). In contrast, *tau ^KO^* tubules displayed enlarged and clustered Lamp1-positive structures, suggesting altered lysosomal organization (Fig. 5H-H’). Quantitative analysis revealed significant changes in lysosomal puncta distribution in *tau ^KO^* tubules compared with controls (Fig. S4C).

Because Rab7-positive late endosomes normally fuse with lysosomes during endosomal maturation, we next examined the spatial relationship between Rab7 and Lamp1. In control tubules, Rab7 puncta frequently overlapped with Lamp1-positive lysosomal compartments, indicating efficient late endosome–lysosome fusion (Fig. 5I–I’’’). In contrast, *tau ^KO^* tubules displayed abnormal accumulation of Rab7-positive vesicles with reduced overlap with Lamp1-positive lysosomes (Fig. 5J–J’’’). Quantification of Rab7–Lamp1 colocalization using Pearson’s correlation coefficient revealed a significant reduction in colocalization in *tau ^KO^* tubules (Fig. 5K), indicating disruption of late endosomal trafficking and lysosomal fusion.

Together, these results demonstrate that loss of dTau disrupts endosomal trafficking and compromises autophagic–lysosomal homeostasis in MTs, leading to defects in autophagosome formation, cargo degradation, and lysosomal organization.

## DISCUSSION

This study reveals a previously unrecognized role for dTau in regulating epithelial growth and tissue architecture through modulation of Notch signaling in the *Drosophila* Malpighian tubules (MTs). While Tau is well known for stabilizing microtubules and supporting axonal transport in neurons (Gotz et al., 2019; Bakota and Brandt, 2024), our findings extend its functional relevance to a non-neuronal epithelial context (Tiwari and Tapadia, 2026), where it contributes to the control of tissue morphology and proliferation.

In the *Drosophila* MTs, Notch signaling plays a critical role during early development by regulating cell fate specification between PCs and SCs and by contributing to proper tubule growth and morphogenesis (Wan et al., 2000; Denholm and Skaer, 2009). Disruption of Notch signaling during MTs development leads to abnormal epithelial organization and altered cell numbers. More broadly, the Notch pathway is a key regulator of epithelial growth, differentiation, and tissue patterning, and its dysregulation frequently results in abnormal tissue expansion and morphogenetic defects (Tricarico and Crovella, 2023; Bray and Bigas, 2025). Consistent with this, *tau ^KO^* tubules display reduced levels of the Notch intracellular domain (NICD), indicating diminished pathway activation. Importantly, reduced Notch activity has been associated with excessive epithelial proliferation and abnormal branching morphogenesis in several epithelial systems, where Notch normally functions to restrict cell proliferation and maintain tissue architecture (Nicolas et al., 2003; Bouras et al. 2008; Surendran et al., 2010). The epithelial hyperplasia and ectopic branching observed in *tau ^KO^* tubules are therefore likely a consequence of impaired Notch signaling. This interpretation is supported by genetic interactions showing that reduction of Notch mimics, and activation of Notch partially rescues, the *tau ^KO^* phenotype.

Microtubules serve as tracks for the movement of endosomal vesicles through motor proteins such as dynein and kinesin (Fourriere et al., 2020; Qu et al., 2025). Disruption of Tau may therefore destabilize these transport pathways, leading to altered endosomal organization, defective receptor trafficking, and reduced Notch activation (Chaudhary et al., 2018; Cason and Holzbaur, 2022; Soppina et al., 2022), consistent with our present results showing the disruption in intracellular trafficking processes in the absence of dTau. Canonical Notch signaling relies on tightly regulated endocytic trafficking, in which receptors undergo internalization, sorting through endosomal compartments, and subsequent proteolytic processing to generate NICD (Yamamoto et al., 2010; Zhou et al., 2022). Disruption of these trafficking steps can significantly affect signaling output. Previous studies have shown that the maturation and recycling of endosomal compartments, mediated by Rab GTPases such as Rab5, Rab7, and Rab11, play critical roles in regulating Notch receptor processing and activity (Vaccar et al., 2008; Fortini, and Bilder, 2009). Consistent with this model, *tau ^KO^* tubules exhibit pronounced alterations in Rab-defined endosomal compartments, including enlargement of Rab5-positive early endosomes, accumulation of Rab7-positive late endosomes, and reduction of Rab11-positive recycling endosomes. These defects suggest impaired progression and recycling within the endocytic pathway, which is essential for productive Notch activation.

Importantly, our proteomic and functional analyses identify the endocytic adaptor Liquid facets (Lqf/Epsin) as a key component affected by Tau loss. Lqf is required for Delta internalization and efficient activation of the Notch pathway (Overstreet et al., 2003; Wang and Struhl, 2004). Reduced Lqf levels in *tau ^KO^* tubules likely impair ligand endocytosis, thereby compromising Notch activation. This probably results in altered localization of the Notch ligand Delta in *tau ^KO^*tubules, with increased cytoplasmic accumulation and membrane enrichment, indicative of defective trafficking. Notably, differences in Delta distribution are also evident in SCs, where the signal appears more diffuse and less organized compared to wild type. Although not directly quantified at the cell-type level, this observation raises the possibility that Tau-dependent trafficking defects may differentially affect specific cell populations within the epithelium.

In addition to changes in classical endosomal compartments, *tau ^KO^* tubules display abnormal accumulation of Ataxin-2–positive ribonucleoprotein (RNP) granules. Ataxin-2 has been implicated in RNA metabolism, stress granule formation, and the regulation of vesicle trafficking, and mutations in Ataxin-2 are associated with several neurodegenerative diseases (Bakthavachalu et al., 2018; Costa et al., 2024). In *Drosophila*, Ataxin-2 has also been reported to influence cytoskeletal organization and endocytic processes (Del et al., 2022). The accumulation of Ataxin-2-positive granules observed in *tau ^KO^* tubules may therefore reflect broader disturbances in cytoplasmic organization and trafficking homeostasis when dTau function is lost. Defects in endosomal transport are also likely to influence downstream degradation pathways that contribute to receptor turnover. Efficient receptor regulation requires coordinated trafficking from endosomes to lysosomes as well as interactions with the autophagic machinery (Langemeyer et al., 2018; von Zastrow and Sorkin, 2021). The accumulation of endosomal markers observed in *tau ^KO^* tubules is consistent with impaired endosomal maturation or turnover, a phenomenon that has been reported in other contexts where vesicular trafficking is disrupted. Such alterations could interfere with the normal processing of Notch receptors, thereby reducing NICD generation and downstream signaling activity.

In neurons, Tau stabilizes microtubules and supports long-distance axonal transport, and disruption of Tau function impairs vesicular movement and synaptic activity (Zhou et al., 2017; Uytterhoeven et al., 2024). The present results suggest that dTau contributes to the organization of cytoskeletal structures that facilitate vesicle transport along microtubules, including in non-neuronal tissues, such as epithelial cells, where microtubule-dependent trafficking is essential for maintaining signaling homeostasis and proper tissue organization.

### Summary

Tau plays a critical role in maintaining epithelial organization and signaling in the *Drosophila* Malpighian tubules. In wild-type tubules, Tau supports proper vesicular trafficking and endosomal organization, enabling efficient Delta internalization and Notch activation, which maintains normal epithelial growth and tubule morphology. In contrast, loss of Tau disrupts vesicle trafficking pathways, leading to altered Rab5, Rab7, and Rab11 endosome distribution and reduced levels of the endocytic adaptor Liquid facets (Epsin). These defects impair Delta trafficking and reduce NICD production despite increased Notch transcript levels. Tau deficiency also disturbs autophagy–lysosomal homeostasis and reduces endosome–lysosome fusion. Together, these defects lead to epithelial hyperplasia, increased tubule diameter, and abnormal branching in MTs.

### Limitation of the study

Although this study identifies a novel role for Tau in regulating epithelial growth and Notch signaling, several limitations should be considered. Our findings are based on analyses in the *Drosophila* Malpighian tubule, and it remains to be determined whether similar mechanisms operate in other epithelial tissues or in vertebrate systems. In addition, while our data suggest that Tau influences Notch activation through effects on endosomal trafficking, the precise molecular mechanisms linking Tau to vesicle transport were not directly examined. Finally, the alterations observed in endosomal compartments and Ataxin-2–positive granules are largely based on static imaging, and future studies using dynamic trafficking assays will be necessary to further define the role of Tau in these processes.

## RESOURCE AVAILABILITY

### Lead contact

Further information and requests for resources and reagents should be directed to and will be fulfilled by the lead contact, Madhu G. Tapadia.

### Materials availability

This study did not generate new unique reagents.

### Data and code availability

- This paper does not report the original code.
- Any additional information required to reanalyze the data reported in this paper is available from the lead contact upon request.

## ACKNOWLEDGEMENTS

We thank Dr. J. A. T. Dow, Prof. Surajit Sarkar, Dr. Bhupendra Shravage and BDSC for sharing their fly stocks. We also acknowledge Prof. K. Vijay Raghavan and DSHB for antibodies. We also thank the Interdisciplinary School of Life Sciences (ISLS), Banaras Hindu University for providing the Confocal facility and Central Discovery Center (CDC), Banaras Hindu University, for providing the HRMS (High Resolution Mass Spectroscopy) facility. This study was funded by Institute of Eminence Scheme, BHU to M.G.T. and UGC fellowships to N. Tiwari.

## AUTHOR CONTRIBUTIONS

Conceptualization, M.G.T. and N. Tiwari; methodology, M.G.T., N. Tiwari and K. Sharma; investigation and analysis, N. Tiwari and K. Sharma; visualization, N. Tiwari and K. Sharma; writing, N. Tiwari; funding acquisition and supervision, M.G.T.

## SUPPLEMENTAL INFORMATION

Document S1. Figures S1–S4

## MATERIAL AND METHODS

### Fly culture and stocks

All *Drosophila melanogaster* stocks, including wild type and mutant lines, were maintained on a standard cornmeal–yeast–agar medium containing 10% yeast, 2% agar, 10% sucrose, 10% autolyzed yeast, 3% nipagin, and 0.3% propionic acid. Cultures were reared at 25 ± 1 °C under a 12 h:12 h light–dark cycle with 50–70% relative humidity. The wild-type strain used in this study was *D. melanogaster* (Oregon-R^+^). The following mutant line was employed: Tau knockout (*tau ^KO^*) mutants (Burnouf et al., 2016; kindly provided by Dr. L. Partridge, Max Planck Institute for Biology of Aging, Cologne, Germany). Additional lines included *UAS-Notch^DN^*, Activated Notch overexpression line (*UAS-NICD*) were generous gifts from Dr. Spyros Artavanis-Tsakonas, *3x Atg8a-mcherry*, from Bhupendra Shravage, Agarkar Research Institute, Pune, *UAS-Lqf* (BL# 57350), *UAS-Lqf RNAi* (BL# 58130), *UAS-Lamp1 GFP* (BL# 42714), *UAS-GFP-Ref(2)P* (BL# 605356). For cell type–specific expression, the principal cell driver *c42 GAL4* and the stellate cell driver *c724 GAL4* were used (kindly provided by Dr. J. A. T. Dow, Institute of Biomedical and Life Sciences, University of Glasgow, UK).

All experiments were performed using Malpighian tubules dissected from wandering third-instar larvae. Virgin females aged three to four days and newly eclosed males were used for all genetic crosses.

### Malpighian Tubule dissection and immunostaining

MTs were extracted from healthy third-instar larvae (118–120 h after hatching) in 1X PBS. Tissue samples were fixed in 4% paraformaldehyde for 20 min at ambient temperature, followed by rinsing with 0.1% PBST (1X PBS with 0.1% Triton X-100). A blocking solution containing 0.1% Triton X-100, 0.1% BSA, 10% FCS, 0.1% deoxycholate, and 0.02% thiomersal was applied to the samples for 2 h at room temperature. The sample tissues were kept in the primary antibodies overnight at 4°C. After this incubation period, the samples were washed three times with 0.1% PBST for 20 min each. An additional 2-hour blocking step was performed before incubation with the secondary antibodies. DAPI (1 mg/ml, Molecular Probe) was used to counterstain the tissues for 15 min at room temperature, followed by further washing with 0.1% PBST. For imaging purposes, the samples were mounted in DABCO antifadant (Sigma, Cat# D27802). The following stains and antibodies were used:

#### Stains

DAPI (1 mg/ml, Thermo Fisher Scientific, Cat# D1306)

#### Primary Antibodies

C594.9B Anti-Delta (DSHB, 1:100, Cat# 528194), C17.9C6 Anti-Notch (DSHB, 1:20, Cat# 528410), Anti-Rab7 (DSHB, 1:20, Cat# 2722471), Anti-Rab5 (CST, 1:1000, Cat# 3547T), Anti-Ataxin2(1:400, provided by Prof. K. Vijay Raghavan), Anti-Atg8 (CST,1:100, Cat# 64459), Anti-Lqf (Abclonal, 1:100, Cat# A20872), Anti-GFP (Invitrogen,1:800, Cat # A10262).

#### Secondary Antibodies

Goat anti-mouse Alexa Fluor 488 (Jackson ImmunoResearch, 1:100), Goat anti-rabbit Alexa Fluor 546 (Invitrogen, 1:200), donkey anti-mouse Alexa Fluor 488 (Invitrogen, 1:200), Goat anti-mouse Alexa Fluor 647 (Invitrogen, 1:200), , Goat anti-rabbit Alexa Fluor 647 (Invitrogen, 1:200), Donkey anti-chicken Alexa Fluor 546 (Invitrogen, 1:200).

### Proteomic Analysis of Malpighian Tubules

#### Protein Extraction and SDS–PAGE

Malpighian tubules were dissected from third instar wandering larvae and collected in cold phosphate-buffered saline (PBS). Tissues were lysed in RIPA buffer supplemented with protease inhibitors using a tissue homogenizer to extract total proteins. The homogenates were centrifuged at 10,000 rpm for 15 min at 4 °C to remove debris, and the supernatants were collected. Proteins were precipitated by adding chilled acetone (1:3 ratio) followed by incubation at −20 °C for 20 h. Samples were then centrifuged to collect protein pellets. Protein concentration was determined using the Bradford assay, and sample quality was assessed by SDS–PAGE (Sharma et al., 2025).

#### In-solution Digestion and LC–MS/MS Analysis

Protein samples were subjected to in-solution tryptic digestion following standard protocols. Briefly, proteins were reduced with dithiothreitol, alkylated with iodoacetamide, and digested with sequencing-grade trypsin overnight at 37 °C. Peptides were desalted using C18 spin columns and analyzed by nano-LC coupled to an Orbitrap Eclipse Tribrid mass spectrometer (Thermo Fisher Scientific) at the Sophisticated Analytical and Technical Help Institute (SATHI), Banaras Hindu University, India. Peptides were separated on a C18 analytical column and analyzed using a data-dependent acquisition method.

#### Protein Identification and Quantification

Raw mass spectrometry data were processed using Proteome Discoverer (v3.1). Protein identification was performed against the *Drosophila melanogaster* protein database, and label-free quantification (LFQ) was used to determine protein abundance. Proteins identified with a false discovery rate (FDR) <1% and adjusted p-value ≤0.05 were considered significant.

Differential expression analysis was performed to identify differentially expressed proteins (DEPs) between control and tau knockout (*tau ^KO^*) MTs. Lists of upregulated and downregulated proteins were compiled and provided as a supplementary Excel dataset. Data visualization and statistical analyses, including bar plots of DEP counts, volcano plots (log_₂_ fold change vs −log_₁₀_ p-value), and protein abundance distribution plots, were generated using GraphPad Prism.

#### Reverse transcription (RT-PCR)

For quantitative RT-PCR analysis, Malpighian tubules were isolated from approximately 50 healthy wandering third-instar larvae for each genotype. Total RNA was extracted using TRIzol reagent (Sigma-Aldrich, Cat# T9424, India) according to the manufacturer’s protocol. The resulting RNA pellet was resuspended in 12 μl of DEPC-treated water, and RNA concentration and purity were assessed spectrophotometrically.

To eliminate potential genomic DNA contamination, 1 μg of total RNA was treated with 1 U of RNase-free DNase I (Thermo Fisher Scientific, Cat# 89836) at 37 °C for 30 minutes. First-strand cDNA synthesis was subsequently performed using DNase-treated RNA as the template following standard reverse transcription procedures.

Quantitative real-time PCR (qRT-PCR) was carried out using the ABI 7500 Real-Time PCR System (Applied Biosystems) with SYBR Green Master Mix (Genetix, Cat# PKG025-A). Each reaction was performed in a final volume of 10 μl containing 5 μl of SYBR Green master mix and 2 pmol/μl of each forward and reverse primer. Relative transcript levels were calculated using the comparative Ct (ΔΔCt) method, with Rp49 used as the internal reference gene for normalization. For each gene, data represent the mean of three independent biological replicates. Primer sequences used in this study are listed below.

*Notch* (Forward) 5’- TCTATTGCCAGTGTACGAAG -3’
*Notch* (Reverse) 5’- ATGTCCTCTGAACAATCCAC-3’
*Lqf* (Forward) 5’- GCAAGGATCAGGGCACCCAT-3’
*Lqf* (Reverse) 5’- ATCGCTGCCGAACCCACTCG -3’
*Rp49* (Forward) 5’- TTGAGAACGCAGCGACCGT-3’
*Rp49* (Reverse) 5’- CGTCTCCTCCAAGAAGCGCAAG-3’

#### Protein extraction and Western blots analysis

MTs from late third-instar larvae (114–118 h after egg laying) of the desired genotypes were dissected in 1× phosphate-buffered saline (PBS). The isolated tissues were transferred to RIPA lysis buffer (Sigma, Cat# R0278) supplemented with a protease inhibitor cocktail (Roche) and homogenized thoroughly. Lysates were centrifuged at 10,000 rpm for 10 minutes at 4 °C, and the clarified supernatant containing soluble proteins was collected. Protein concentration was measured using the Bradford assay.For each sample, 30 µg of total protein was combined with SDS sample buffer (100 mM Tris–HCl, pH 6.8; 4% SDS; 20% glycerol; 100 mM DTT; 2% β-mercaptoethanol; and 0.2% bromophenol blue) and denatured by heating at 95 °C for 5 minutes.

Proteins were resolved on 10% SDS–polyacrylamide gels at 100 V for approximately 1 hour and subsequently transferred onto PVDF membranes (Millipore) using a Bio-Rad Mini-Protean transfer apparatus at 120 mA for 2 hours at 4 °C. Membranes were blocked for 1 hour at room temperature with 4% BSA (SRL, Cat# 83803) prepared in TBST (TBS containing 0.1% Tween-20) to prevent nonspecific binding. The membranes were then incubated overnight at 4 °C with primary antibodies diluted in blocking buffer: anti-Notch C17.9C6 (DSHB, 1:500, Cat# 528410), anti-β-tubulin E7 (DSHB, 1:1000, Cat# 528499), anti-Lqf (ABclonal, 1:1000, Cat# A20872) and anti-Delta C594.9B (DSHB, 1:1000, Cat# 528194). Following primary antibody incubation, membranes were washed three times with 0.1% TBST for 15 minutes each. The blots were then incubated with HRP-conjugated anti-mouse (Invitrogen, 1:1000, Cat# 31430) and anti-rabbit (Invitrogen, 1:1000, Cat# 32460) secondary antibody for 2 hours at room temperature.

After three additional washes with TBST, protein bands were detected using an enhanced chemiluminescence (ECL) substrate (Bio-Rad) and visualized using a ChemiDoc imaging system (Amersham 680).

#### Microscopy and image processing

All imaging was performed using Zeiss LSM 900 confocal laser scanning microscopes operated with ZEN software (version 3.4). Z-stack images were collected with an optical section interval of 2.0 µm unless otherwise specified in the relevant figure legends. Appropriate laser lines and emission filters were selected for each fluorophore to ensure optimal signal detection. Image processing and quantitative analyses were carried out using ImageJ software (NIH, USA; https://imagej.nih.gov/ij/). Final figures were prepared using Adobe Photoshop 2021 (version 22.4.2). Schematic diagrams and graphical illustrations were generated using BioRender and Microsoft PowerPoint.

#### Image quantification and statistical analysis

Fluorescence image quantification was performed using ImageJ (Fiji, NIH). For intensity measurements, a region of interest (ROI) encompassing the entire MTs was selected, and the mean fluorescence intensity was calculated from 10 independent optical sections per sample to ensure consistency across genotypes.

For puncta-based analyses, the number of fluorescent puncta within a defined ROI was quantified using threshold-based particle analysis in ImageJ. The total number of puncta per ROI was recorded, and puncta density was calculated relative to the analyzed area. In addition, the average size of puncta per cell was measured by determining the area of individual particles detected within the ROI.

Colocalization between Rab7 and Lamp1 signals was quantified using Pearson’s correlation coefficient (PCC) calculated with the *JACoP* plugin in ImageJ. Pearson’s coefficient values range from −1 to +1, where values close to +1 indicate strong positive colocalization, values near 0 indicate random distribution, and values approaching −1 indicate negative correlation.

All quantitative data were collected from at least three independent biological replicates. Graphs and statistical analyses were generated using GraphPad Prism (version 7.0). Comparisons between two groups were performed using a two-tailed unpaired Student’s *t*-test, while comparisons involving multiple groups were analyzed using one-way ANOVA followed by Tukey’s post hoc test. Data are presented as mean ± standard error of the mean (SEM), and differences with *p* < 0.05 were considered statistically significant.

