## Supplementary Data for "Tau regulates epithelial morphogenesis through vesicle trafficking–dependent Notch activation"

**Supplementary Information:**

**Supplemental Figure and Legend:**

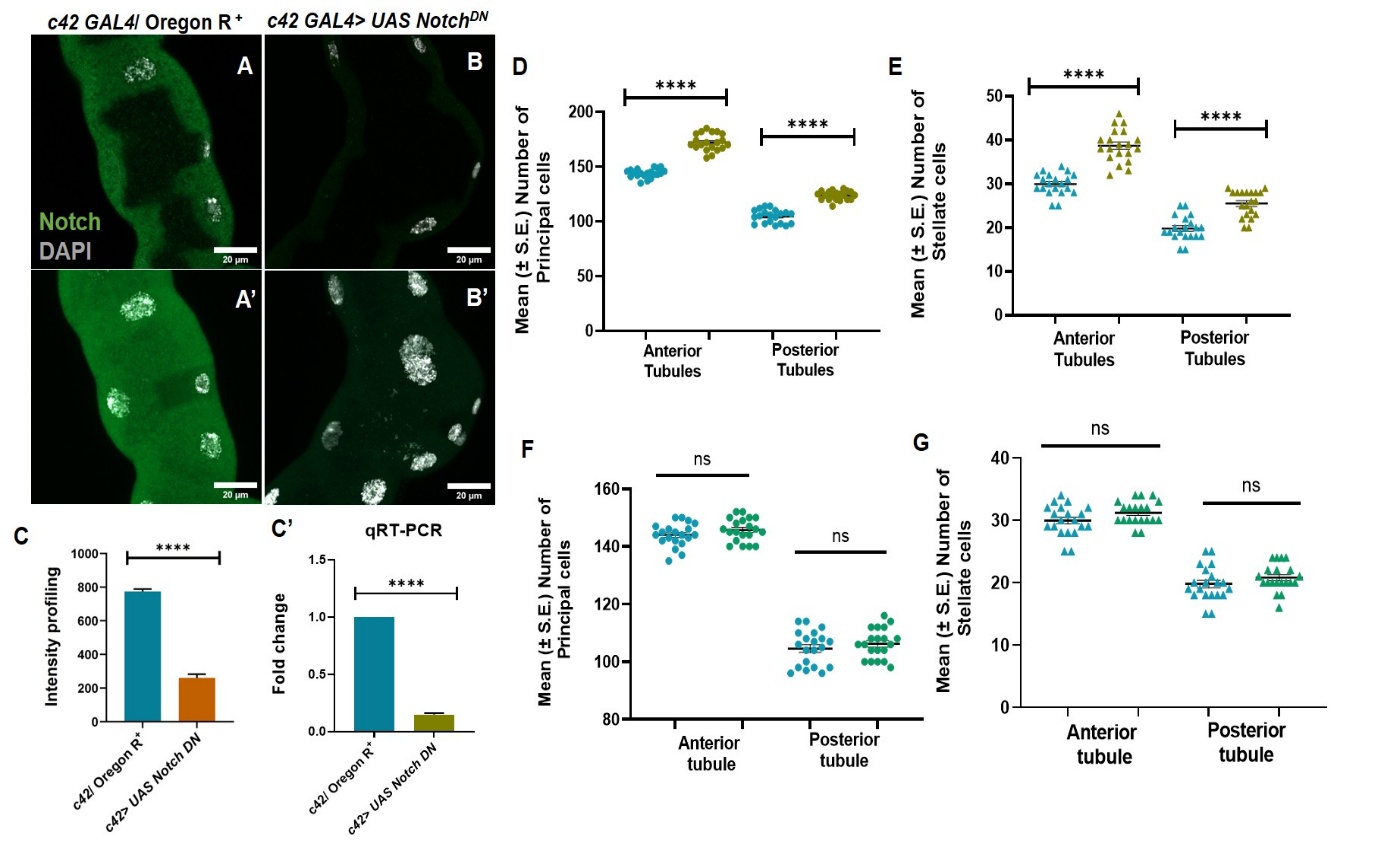

**Figure S1: Loss of Notch phenocopies Tau loss in MTs.**

#### (A-A’) Wild-type tubules showing NICD (green) with DAPI (grey) staining in 3^rd^ instar MTs.

#### (B-B’) NICD immunostaining (green) in *c42 GAL4> UAS Notch^DN^* 3^rd^ instar MTs.

**(C)** Quantification of NICD fluorescence intensity in MTs (n=10).

**(C’)** RT–PCR analysis of Notch transcript levels in tubules (n=3).

**(D-E)** Scatter dot plot showing the mean (±SE) number of principal cells (PCs) and stellate cells (SCs) (n = 10 pairs of MTs per genotype) in control and *c42 GAL4> UAS Notch^DN^* condition (n=20).

**(F-G)** Scatter dot plot showing the mean (±SE) number of principal cells (PCs) and stellate cells (SCs) (n = 10 pairs of MTs per genotype) in control and *c42-GAL4 > UAS-NICD;* *tau ^KO^* condition (n=20).

All the Malpighian tubule images shown are from the wandering third-instar larvae. Scale bars represent 20 μm. All images are maximum intensity projections of all the sections except (A’ and B’) which are single section images. DAPI (grey)-stained nuclei. Statistical analysis was done using unpaired t-test, ****p < 0.0001. Error bars, mean ± SE. All images represent 3 or more independent biological replicates.

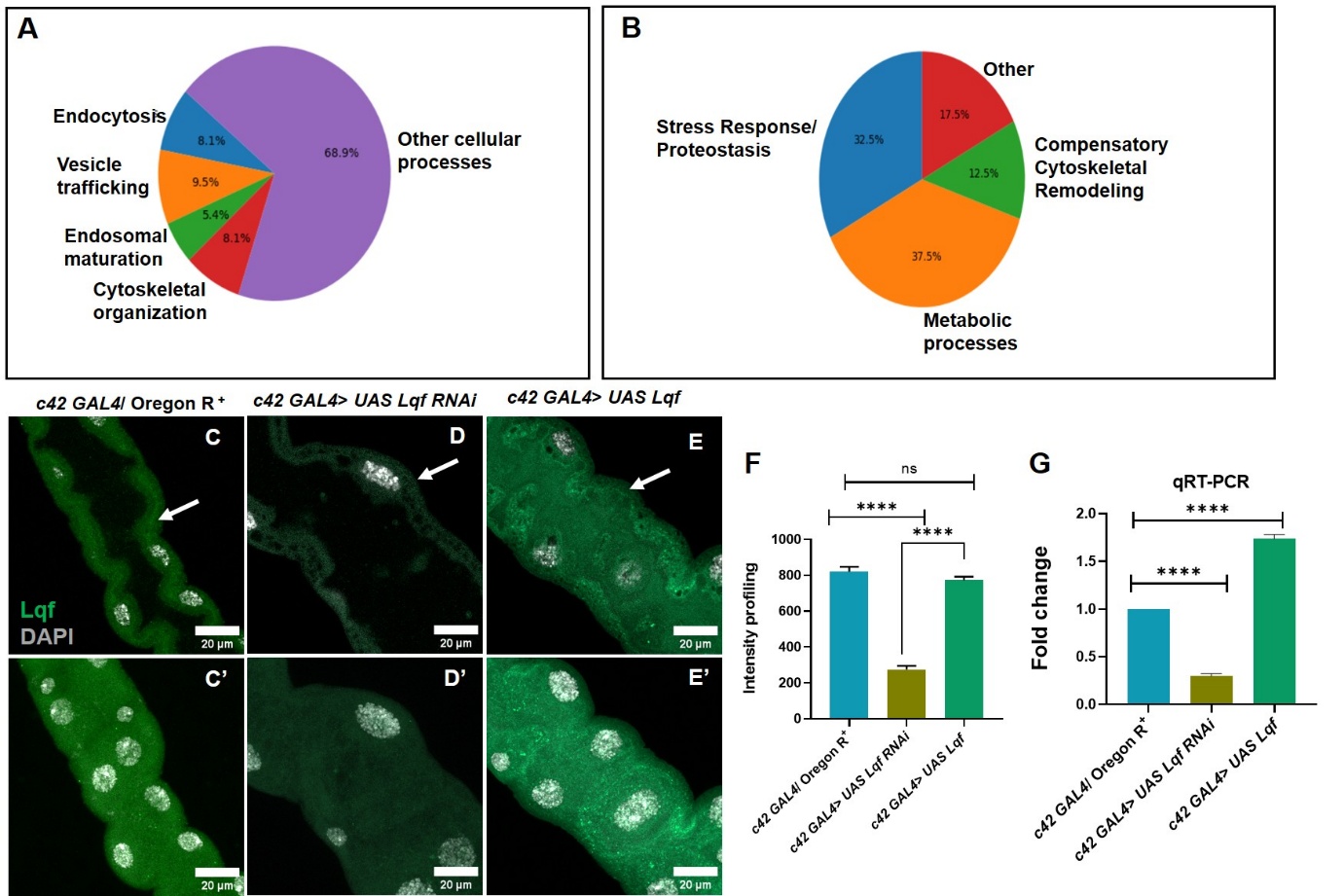

### **Figure S2. Functional classification of proteomic changes and effect of Lqf manipulation on Notch signaling**

**(A)** Pie chart showing functional categorization of proteins significantly downregulated in *tau ^KO^* MTs.

**(B)** Pie chart showing functional categorization of proteins significantly upregulated in *tau ^KO^* MTs.

**(C-C’)** Immunostainig of NICD (green) in control MTs.

**(D–D″)** Immunostaining of NICD (green) in MTs following RNAi-mediated knockdown of Lqf using *c42-GAL4*.

**(E–E″)** Immunostaining of NICD (green) in MTs overexpressing Lqf (*c42-GAL4 > UAS-Lqf*).

**(F)** Quantification of NICD fluorescence intensity in MTs (n=10) .

**(G)** RT–PCR analysis of Lqf transcript levels in different genetic background (n=3).

All the Malpighian tubule images shown are from the wandering third-instar larvae. Scale bars represent 20 μm. All images are maximum intensity projections of all the sections except (C, D and E) which are single section images. DAPI (grey)-stained nuclei. Statistical analysis was done using one-way ANOVA followed by Tukey’s multiple comparison test, ****p < 0.0001. Error bars, mean ± SE. All images represent 3 or more independent biological replicates.

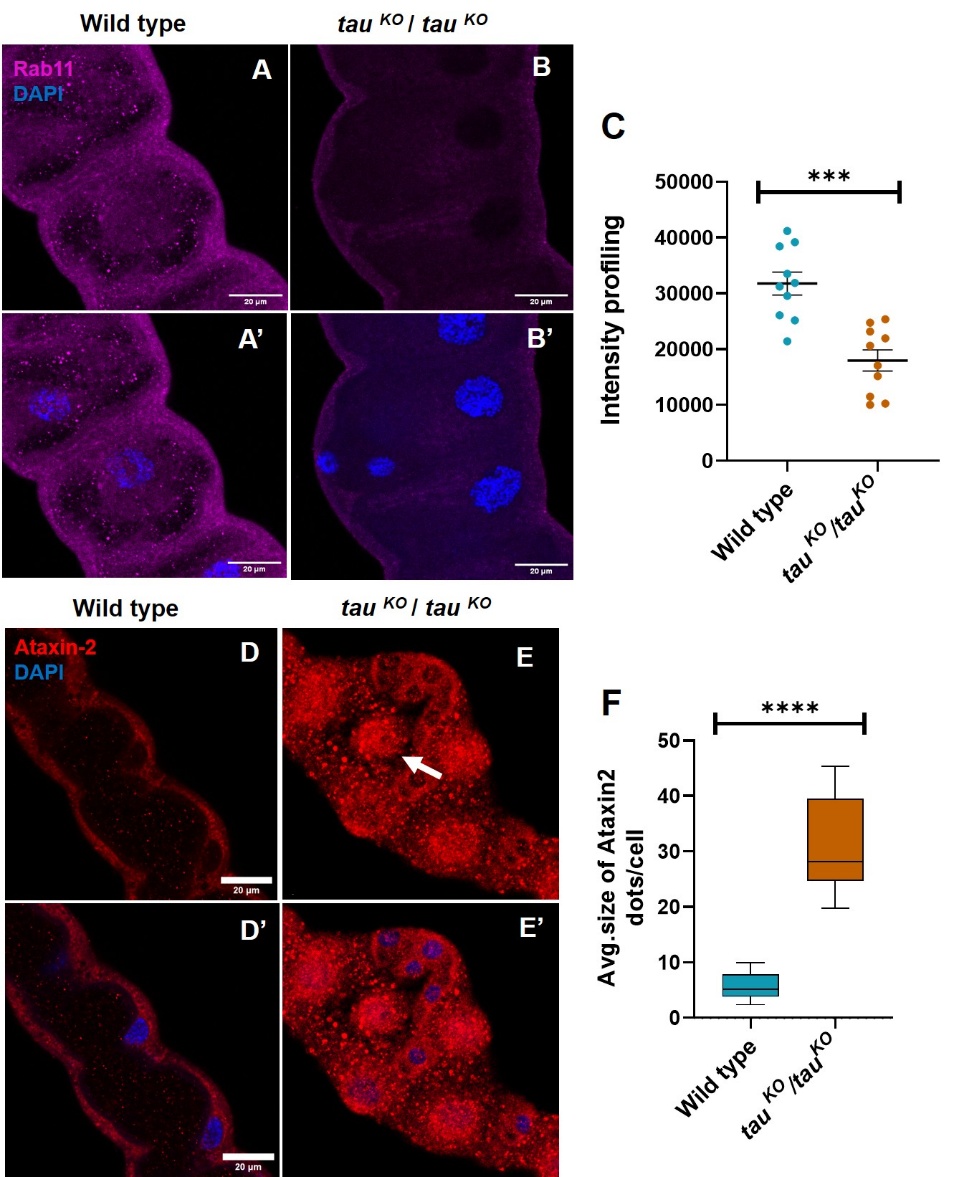

**FigureS3: Tau loss disrupts Rab11-positive recycling endosomes and promotes Ataxin-2 aggregation in Malpighian tubules.**

**(A–A′)** Representative confocal images showing the localization of Rab11 (magenta) in wild-type MTs.

**(B–B′)** Representative confocal images showing the localization of Rab11 (magenta) in *tau ^KO^* MTs.

**(C)** Quantification of Rab11 fluorescence intensity in wild-type and *tau ^KO^* tubules (n=10).

**(D–D′)** Confocal images showing staining of Ataxin2 (red) in wild-type tubules.

**(E–E′)** Confocal images showing staining of Ataxin2 (red) in *tau ^KO^* tubules.

**(F)** Quantification of the average size of Ataxin-2–positive puncta per cell shows a significant increase in *tau ^KO^* tubules (n=10).

All the Malpighian tubule images shown are from the wandering third-instar larvae. Scale bars represent 20 μm. All sare single section. DAPI (blue)-stained nuclei. Statistical analysis was done using unpaired t-test, ***p<0.001 and ****p < 0.0001. Error bars, mean ± SE. All images represent 3 or more independent biological replicates.

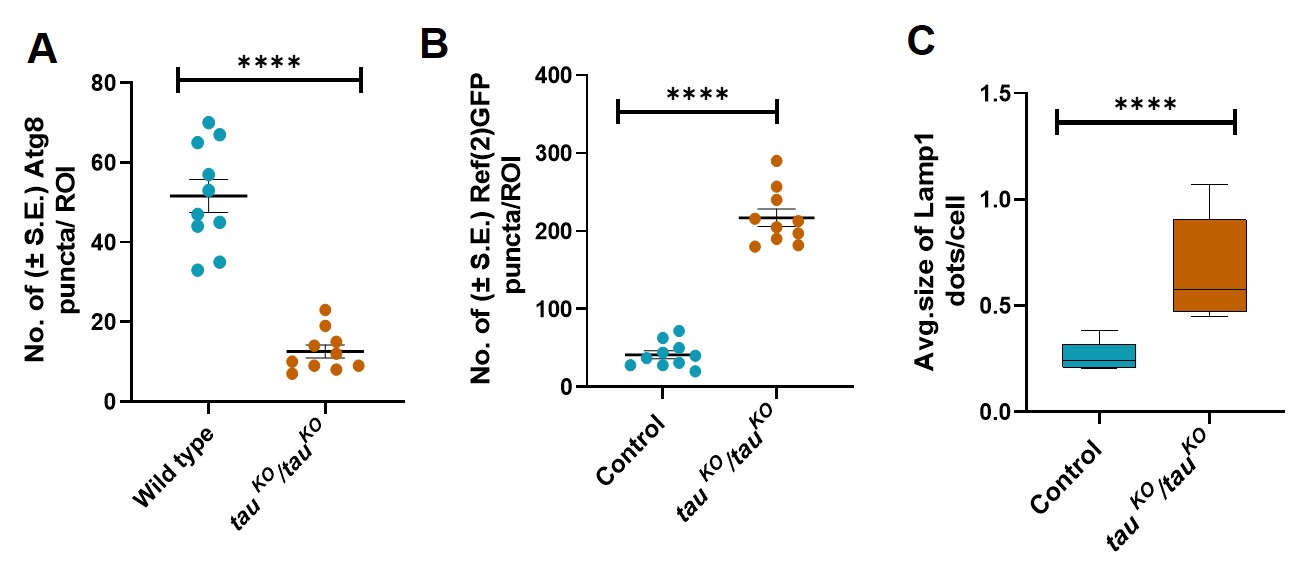

##### **Figure S4. Quantification of autophagic and lysosomal markers in Malpighian tubules.**

**(A)** Quantification of **Atg8 puncta** in control and *tau ^KO^* MTs (n=10).

**(B)** Quantification of **Ref(2)P puncta** in control and *tau ^KO^* tubules (n=10).

(C) Quantification of **Lamp1-positive lysosomal puncta size per cell** in control and *tau ^KO^* tubules (n=10).

Data represent mean ± SEM from multiple tubules analyzed per genotype. Statistical significance was determined using an unpaired Student’s t-test, ****p < 0.0001.
